# Pathological and non-pathological aging, SAMP8 and SAMR1. What do hippocampal neuronal populations tell us?

**DOI:** 10.1101/598599

**Authors:** MJ Lagartos-Donate, J Gonzáles-Fuentes, P Marcos-Rabal, R Insausti, MM Arroyo-Jiménez

**Affiliations:** Kavli Institute for System Neuroscience Centre for Neural Cumputation; Universidad de Castilla-La Mancha, Facultad de Farmacia, Albacete, Spain; Universidad de Castilla-La Mancha, Facultad de Medicina, Albacete, Spain

**Keywords:** aging, SAMP8, interneuron, parvalbumin, somatostatin, calretinin

## Abstract

Alterations of cognitive processes and memory are one of the most important manifestations related to aging. However, not all memory types are affected in the same way. Learning and spatial memory are susceptible to these changes. The hippocampus represents the anatomical substrate of this type of memory, affected by structural and functional alterations along the normal aging and neurological diseases such as Alzheimer’s disease, Parkinson’s, schizophrenia and epilepsy. Some of the alterations related to aging are associated with alterations in the hippocampal interneuron populations and with an increase in excitability in the hippocampal circuit.

In order to understand better the underlying processes in normal and pathological aging mechanisms, a murine model `(Senescence-Accelerated Mouse Prone, SAMP8)` and its respective controls `(Senescence-Accelerated Mouse Resistant, SAMR1)` have been used. While SAMP8 is a naturally occurring mouse line that displays a phenotype of accelerated aging with learning and memory impairment and these changes of learning and memory might be linked to some alterations in neuronal populations of the hippocampus. Thus, we analyzed the distribution and density of PV, CR and STT interneurons in the hippocampus of young and old mice as well as possible morphological and cholinergic changes in hippocampal formation. Comparing SAMR1 and SAMP8 we did not find any neural population that was specifically affected by aging in both groups. Interestingly, CR immunoreactivity and STT immunoreactivity showed changes in SAMP8 mice when they were compared to their controls. In SAMP8 CR+ and STT+ neurons decreased significantly along aging which suggests that CR and STT interneurons play a more important role than PV neurons in the pathological aging of the brain. In the case of SAMP8 mice the neural changes might be related to changes of the cholinergic system that might be affecting the wiring into the hippocampus formation through the perforant pathway. Further studies of this local circuitry will help to comprehend better how different inputs into these neural populations of the hippocampus could be affecting the development of neurodegerative diseases.

## INTRODUCTION

Aging and aging-related neurodegenerative disorders like Alzheimer’s disease (AD) are one of the major health challenges of the modern societies. In order to improve the quality of life of our society, it is crucial to understand the relative contribution of multiple anatomo-functional and physiological changes that often precede the appearance of clinical signs of cognitive decline.

One of the most remarkable declines and the earliest manifestation of cognitive senescence is impaired memory. However, not all types of memory are equally susceptible to aging. While long-term retention of previously acquired skills and procedures show more resistance to age-related damage (Foster, Defazio et al. 2012), learning and memory, which encompass knowledge of location within an environmental context, are particularly vulnerable to age related decline in humans and rodents (Davis, Markowska et al. 1993, Lindner 1997, Foster, Sharrow et al. 2003, Driscoll, Howard et al. 2006, Bizon, LaSarge et al. 2009, Foster, Defazio et al. 2012).

We know that anatomical substrates for episodic and spatial memory are similar in humans and animal models of aging and neurodegenerative diseases. In that line, hippocampus, a particularly vulnerable structure which is known to undergo structural (for example decrease of volume) and functional alterations during normal and pathological age-related process (Burke and Barnes 2006, Wilson, Tang et al. 2006), is included among these relevant brain substrates (Nadel and Moscovitch 1997, Nadel, Samsonovich et al. 2000). Some of these changes seem to be connected with perturbations in inhibitory networks. Many morphological studies provide evidence for age related changes in inhibitory interneurons of the hippocampus, which include loss of subgroups of GABAergic inhibitory interneurons (Vela, Gutierrez et al. 2003, Shi, Argenta et al. 2004, Potier, Jouvenceau et al. 2006, Kuruba, Hattiangady et al. 2011, Stanley, Fadel et al. 2012). However, the extent and consequences of these alterations are largely unknown.

Numerous studies have shown that the density of GABAergic interneurons declines because of age in area CA1 of the hippocampus (Shetty and Turner 1998, Shi, Argenta et al. 2004). Nevertheless, what it is not so clear is which interneurons populations are vulnerable to normal aging process, and what the difference with a pathological neuronal loosing is. Interneuron populations such as those containing somatostatin (SST), calretinin (CR), or parvalbumin (PV) decrease with aging process, however there are contradictory data in the literature. For instance, the fate of PV+ interneurons during aging is unclear, PV+ cells was observed both unchanged (Vela, Gutierrez et al. 2003, Potier, Jouvenceau et al. 2006) and decreased (Shetty and Turner 1998, Lee, Hwang et al. 2008) in aged animals. On the other hand, a type of SST+ interneurons, known as oriens-lacunosum moleculare (O-LM) cells, are particularly vulnerable to aging(Cardoso, Silva et al. 2014).

However, in AD animal models, like double transgenic PS1/AβPP mice and Tg2576 mice, a loss of hippocampal interneurons is observed in a recurrent manner. Loss of CR+ cells is shown in double transgenic PS1/AβPP mice. As early as 4 months of age, coinciding with the onset of extra-cellular Aβ pathology, the cell number of CR+ interneurons is significantly reduced in CA1 (Baglietto-Vargas, Moreno-Gonzalez et al. 2010). Similarly, 5.5 months-old Tg2576 mice display a significant decrease in SST+ cells, accompanied by a loss of spine density of basal proximal dendrites of the principal cells in the stratum oriens of CA1 region (Perez-Cruz, Nolte et al. 2011). Thus, age-related impairments in learning and memory are associated with decreased number of GABAergic interneurons and increased excitability of hippocampal network (Stanley, Fadel et al. 2012).

In order to have a better understanding of the underlying mechanisms of both normal and pathological aging, we have analyzed three interneuron populations, containing CR, PV and SST in the hippocampus of senescence-accelerated mouse resistance-1 (SAMR1) and senescence-accelerated mouse prone-8 (SAMP8) murine models which show normal and accelerated senescence-process respectively.

The SAMP8 and its control, SAMR1, were established through selective inbreeding of the AKR/J strain based on phenotypic variations of accelerated aging (Takeda, Hosokawa et al. 1981, Hosono, Hanada et al. 1997, Wyss, Kadish et al. 2003, Luo, Li et al. 2009). SAMP8 mice display age-associated impairments in the central nervous system (CNS) function, including marked deficits in learning and memory (Flood and Morley 1993, Miyamoto 1997, Flood and Morley 1998), altered emotional status (Miyamoto, Kiyota et al. 1992, Markowska, Spangler et al. 1998), and abnormality of circadian rhythm ((Miyamoto, Kiyota et al. 1986, Colas, Cespuglio et al. 2005). Furthermore, neuropathological and neurochemical studies have shown changes in SAMP8 brains, such as increased oxidative stress, occurrence of Aβ-immunoreactive deposits and changes in the cholinergic system(Del Valle, Duran-Vilaregut et al. 2010).

Our aim was analyze morphological brains features of these models that might explain some of the observed behavior. For this purpose, we quantified the distribution of PV+, CR+ and SST+ neurons in the hippocampus of young-adults, adults and old mice and we compared the results obtained in the two animal models.

## MATERIAL AND METHODS

### Mouse strain

28 mice were used in this study: 5, 6months-old SAMR1 mice; 5, 6 months-old SAMP8 mice; 4, 12 months-old SAMR1 mice; 4, 12 months-old SAMP8 mice; 5, 16months-old SAMR1 mice and 5, 16 months-old SAMP8 mice.They were separated in several groups according to age and genotype. SAMR1 and SAMP8 mice at the age of 6, 12 and 16 months old were used for this study. All animals were housed under standard conditions for humidity, light and temperature and provided free access to food and tap water. All procedures were approved by the Ethical Committee for Animal Research of Castilla-La Mancha University (PR-2016-05-13) according to Spanish law (RD 1201/2005) and European directive 2010/63/EU).

### Tissue processing

After deep anesthesia with a combined dose of ketamine hydrochloride (75 mg/Kg) and xylacine (10 mg/Kg), mice were perfused transcardially with 0.9% saline solution followed by 4% paraformaldehyde. Brains were dissected from the skull and post-fixed overnight in the same fixative at 4<u>°</u>C.

The brains were cryoprotected by immersion in 30% sucrose solution. Afterwards, the brains were cut at 50µm on a sliding microtome (Microm, Heidelberg, Germany) coupled to a freezing unit into coronal (Left hemisphere) and sagittal sections (Right hemisphere). One-in-five sections was immediately mounted on a gelatin-coated slide for Nissl staining with 0.25% thionin, adjacent sections were used for immunohistochemistry for ChAT SST, CR and PV.

After sections intended to Nissl staining were air dried and placed directly into 1:1 alcohol/chloroform for an hour and then rehydrate through 100%, 96%, 80% and 50% ethanol solutions. Afterwards tissue was placed into 0.25% Thionin solutions for 10 seconds and dehydrated using 80%, 96%, 100% ethanol solutions and finally left into Xylol solutions to be covered with DPX.

### Immunohistochemistry (IHC)

Sections were extensively rinsed in Tris-buffered saline (TBS. 0.01M, pH 7,6), followed by the elimination of endogenous peroxidase using methanol and hydrogen peroxide (2:1). After washes in TBS, sections pre-incubated with a TBS solution containing 1% normal horse serum and 0.3% Triton X-100. This solution was also used for pre-incubation rinses and to dilute the primary and secondary antibodies as well as streptavidin coupled with peroxidase used in the second immunohistochemical reaction.

Sections were incubated separately with three primary antibodies for 24 - 48 hours at 4<u>°</u>C, a goat anti-ChAT (ref. AB144P, Chemicon), a goat anti-SST (D-20. ref. sc-7819, Santa Cruz), a sheet anti-CR (ref. 7699/3H Swant) and a rabbit anti-PV (ref. PV 25, Swant). All the primary antibodies were incubated at 1:1000 dilutions except AChE which were diluted 1:500. After being washed in TBS, sections were incubated for 90 min at room-temperature in the appropriate biotinylated secondary antibody (anti-goat for ChAT and SST, anti-rabbit for PV and anti-Sheep for CR. Working dilution 1:1000), thereafter with peroxidase-conjugated streptavidin for a same tame and dilution than secondary antibody (Jackson Laboratories). The colour reactions was then developed using the 3,3’-diaminobenzidine (DAB) method (DAB. 0.25% in TBS) and 0.01% hydrogen peroxide. Finally, sections were dehydrated and coverslipped with DPX.

For double immunofluorescence analysis we carried out similar protocol, sections were incubated for 48h at 4<u>°</u>C with anti-SST and anti-CR primary antibodies (dilution 1:1000) at the same time. After washing with TBS, the sections were incubated for 90 min at room-temperature with secondary antibodies. Alexa-488 anti-goat (ref. A11057, Invitrigen. Dilution 1:1000) for SST detection, whereas an anti-rabbit biotinylated secondary antibody (ref. Ab6828, Abcam. Dilution 1:1000) and streptavidin coupled to Alexa-568 (ref. A32354, Invitrigen. Dilution 1:1000) for CR detection. Sections were coverslipped with Vectashield (Vector).

### Antibodies specificity

Controls for single immunolabeling were carried out by replacing the primary antibody with blocking solution, or pre-immune goat or horse serum. In addition, negative controls were included in all immunohistochemical experiments by omitting the secondary antibody or replacing it with an inappropriate secondary antibody. Specificity of the immunolabeling was checked on peripheral tissues. Controls for double immunolabeling were carried out by omitting either anti-SST or anti-CR antibody during the first incubation bath and then incubating the sections with all secondary immunoreagents. No immunolabeling was detected.

### Quantification

The distribution and density (immnureactive cells/mm^2^) of SST, CR and PV immunoreactive neurons were charted with a computerized stage (AccuStage, Minnesota Datametrics MD-5 Digitizer, software MDPLOT v5). The density of cells was measured in Stratum oriens, Stratum piramidale, Stratum Radiatum and Stratum lacunosum-moleculare, as well as in the whole surface of the hippocampus which include Dentate Gyrus (DG), CA3, CA2 and CA1. Six animals of each group of age belonging to SAMP8, SAMR1, Pol μ +/+ and Pol μ -/- were used and three sections per animal were analyzed. Mean values ± SD from all the experimental data, are presented in the figures.

The images were obtained by a light microscope (Nikon Eclipse 80-i) and were processed digitally for the adjustment of brightness and contrast by using Adobe Photoshop CS 8.0.1 software.

A confocal microscope was used to study co-localizacion of SST and CR. The images were analysed using ImageJ and ZEN software (confocal microscope, LSM 710, Zeiss). The percentage co-localization was calculated using the profile and ortho tools of the ZEN software.

### Volumetric analysis

Nissl-stained sections were used to estimate the CA1 strata and hippocampal volume. Hippocampal and sub-hippocampal volumes were measured using the Cavalieri estimator using the software ImageJ. Images were captured using a Nikon Eclipse 80i light microscope with a 10× objective coupled to a Nikon color camera. A series of contiguous images of the Nissl stained samples were obtained, every 5th sections (250 µm thickness between sections) was sampled. Coronal and sagittal sections, obtained from difference hemisphere of the same animal, were used to measure the hippocampus. The first section selected at random following the appearance of the hippocampus. The sagittal measure of the hippocampus starts 0.96 from Bregma and 2.40 lateral to the midline, whereas the coronal measure from −1.22 to −2.54 relative to Bregma (Howard and Reed 2004). Volume was calculated using the formula V = D • ∑A, where D is the length between sections (250 µm), and ∑A is the area of the hippocampus in the sections.

### Densitometry

Densitometry was carried out from IHC against AChE and SST. Optical density (OD) was obtained from 5 sections per animal. Stratum oriens and whole hippocampus were measured from AChE-labeled sections, whereas only hippocampus was measured from SST-labeled sections. The software ImageJ was used to measure the OD from images of the IHC. The Stratum oriens measurements were obtained from four 50 µm^2^ racks. The white matter was taken like background measure.

### Statistical Analysis

Statistical analysis was performed using SPSS for Windows software (version 19.0). Cell density data were subjected to repeated measures analysis of variance (ANOVA) with two between-subjects factors (age and genotype) and one within-subject factor (section). Separate repeated measures ANOVAs with one between-subjects factor (age or genotype) were performed, when needed, for each level of the other factor (genotype or age) in order to elucidate specific differences. When statistical significance was revealed, post hoc tests were performed (Tukey HSD). Differences were considered statistically significant when p < 0.05.

## RESULTS

We first describe the volumetric analysis of the hippocampus in both SAMR1 and SAMP8 mice. Next, we describe the immunohistochemistry of PV, CR and SST expressing neurons in the whole CA1 area and in it subfields. Finally, we analyzed the double staining SST/CR in CA1 of both SAMR1 and SAMP8 mice. Fluorescence and DAB immunohistochemistry techniques have been conducted for all the quantification analysis of PV, CR, and SST neurons. And fluorescence and DAB data were complementary and they support each other, finding significance differences in the same cases.

### VOLUMETRIC HIPPOCAMPAL STUDY

Coronal sections were used to assess changes in hippocampal volume along anterior-posterior axis. The hippocampal volume measurements were taken at three different age points (6, 12 and 16months). The dorsal hippocampus of SAMP8 showed similar volumes as in SAMR1 mice (6 months - SAMR1: 2.60 ± 0.25; SAMP8: 2.79 ± 0.29 mm3; 12 months: SAMR1: 2.67 ± 0.45; SAMP8: 3.03 ± 0.41 mm^3^; 16 months: SAMR1: 2.88 ± 0.37; SAMP8: 2.51 ± 0.20 mm^3^). Therefore, these data did not show statistical differences between the two groups (p-value < 0.05) (supplementary figure 1).

#### 1. NEUROCHEMICAL ANALYSIS

##### A. Parvalbumin interneurons

No qualitative differences in the pattern of distribution of PV+ cells were found between SAMR1 and SAMP8 mice. PV+ neurons were mainly distributed in *stratum oriens* and *stratum pyramidal* of the CA1 field, and a minor number was found in *stratum radiatum.* The study of PV+ cells in CA1 did not show statistical differences between SAMR1 and SAMP8 groups at any of the analyzed ages. At 6 and 12 months-old mice density of PV+ neurons was similar in SAMR1 and SAMP8 mice (6 months - SAMR1: 29.17 ± 10.17; SAMP8: 28.22 ± 6.51 cells/mm^2^. 12 months - SAMR1: 38.92 ± 12.96; SAMP8: 39.70 ± 10.04 cells/mm^2^). Likewise, there was no significant increase between 16 months-aged SAMR1 and 16 months-aged SAMP8 mice (SAMR1: 33.62 ± 9.71; SAMP8: 39.16 ± 8.41 cells/mm^2^) (supplementary figure 2). The density of PV+ neurons did not change significantly in SAMR1, and the same happened in SAMP8 group between 6 months-old mice and 16 months-old mice.

##### B. Calretinin interneurons

CR+ neurons had the same pattern of distribution in SAMP8 and SAMR1 mice at the three points of age analyzed. Our results showed perikarya of CR+ interneurons were mainly found in *stratum piramidale* of CA1, although CR+ interneurons were found in *stratum oriens* and *radiatum* as well.

Quantification of CR+ interneurons revealed no changes of density during aging in CA1 region of SAMR1 mice. On the contrary, it was observed a significantly fall of the density of CR+ cells (p-value= 0.01) between 6 months and 16 months-old SAMP8 mice. Between 6 months and 12 months-old SAMP8 mice the CR+ density decreased up to 20% (and p-value = 0.032), but no significant changes were observed between 12 months-old and 16 months-old SAMP8 mice(figure 1A).

**Figure 1.**
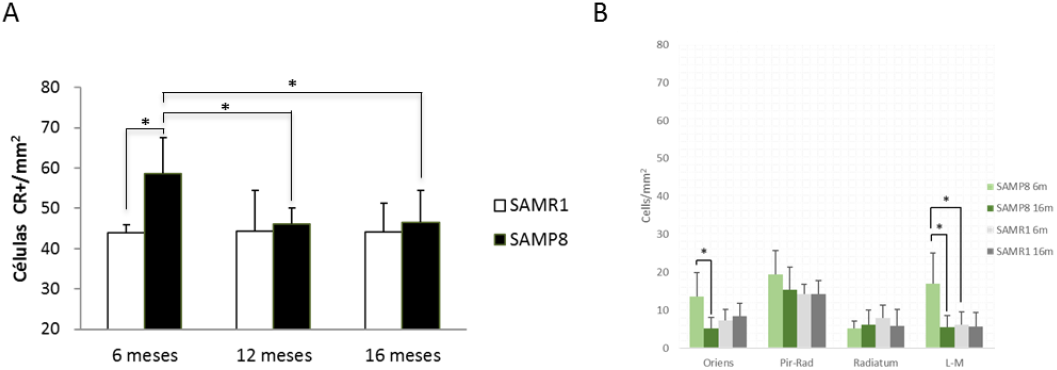
CR immunoreactive neurons in CA1. A) Count of CR+ neurons per mm^2^ in wholfe CA1 field. Measurements in SAMR1 and SAMP8 mice at three different points of age: 6 months-old, 12 months-old and 16 months-old. B) By using ImageJ software and confocal picture we measured the number of CR neurons per mm^3^. The graph shows the density of CR neurons in each of the CA1 subfields. *; p-value ≤0.05.

Comparing SAMP8 and SAMR1 groups, younger ages show significant differences. Interestingly, the density of CR+ interneurons in 6 months-old SAMP8 mice group was up to 31% largest than in 6 months-old SAMR1 mice (SAMR1: 44.09 ± 4.6; SAMP8: 58.16 ± 15.67 cells/mm^2^; p-valor: 0.027). Nevertheless, no significant differences were found between 12 months-old SAMR1 and 12 months-old SAMP8 (SAMR1: 44.38 ± 12.69; SAMP8: 46.06 ± 9.59 cells/mm2), and the same happened between SAMP1 and SAMP8 of 16 months-old (SAMR1: 42.11 ± 7.85; SAMP8: 45.21 ± 10.78 cells/mm^2^) (figure 1A).

Additionally we tried to locate in which of the subfields of CA1 the CR+ population density was changed. The density of CR+ neurons was independently measured in *stratum oriens, pyramidal, radiatum* and *lacunosum-moleculare*. In SAMR1, the highest density of the CR+ neurons was found in *stratum pyramidale*. The values of density and the distribution of CR+ neurons in CA1 field did not change between the young and the old SAMR1 animals. Nevertheless, the distribution of the CR+ neurons was modified in the younger SAMP8 mice. Contrary to 6 months-old SAMR1 mice, the density was higher in the *stratum oriens, pyramidal* and *lacunosum-moleculare* of 6months-old SAMP8 mice while the number of CR+ neurons/mm^2^ in *stratum radiatum* was similar to SAMR1. Although there were 15-20 CR+ neurons/mm^2^ in the previously mentioned subfields at the age of 6 months old, in 16 months-old SAMP8 the density of CR+ neurons of the *stratum oriens* and *lacunosum-moleculare* decreased significant values and reach similar values as SAMR1. Because of this change of CR population in SAMP8, we found significant differences between younger SAMR1 and younger SAMP8 mice in *stratum oriens* and *lacunosum-moleculare* (figure1B).

##### C. Somatostatin interneurons

SST+ cells were mainly found close to the *alveus* in the *stratum-oriens* of CA1 of both SAMR1 and SAMP8. The pattern of distribution of SST+ interneurons was the same in all analyzed groups (6 months-old, 12 months-old, 16 months-old SAMR1 and SAMP8).

A high density of SST+ neuropil was observed in the *stratum lacunosum-moleculare* as well.

The quantitative analysis of interneurons showed no significant differences in SAMR1 during aging, so the number of SST+ cells per area was similar in SAMR1 mice at 6, 12, and 16 months old. However, a striking difference was observed in SAMP8 group between the youngest and elder animals. In that way, SST+ cells decreased a 42% from 6 months to 12 months-old SAMP8 mice (p-value < 0.001) and a 48% from 6 months to 16 months-old SAMP8 mice (p-value < 0.001). Nevertheless, there was no statistical difference between 12 months-old and 16 months-old SAMP8 mice (figure 2A).

**Figure 2.**
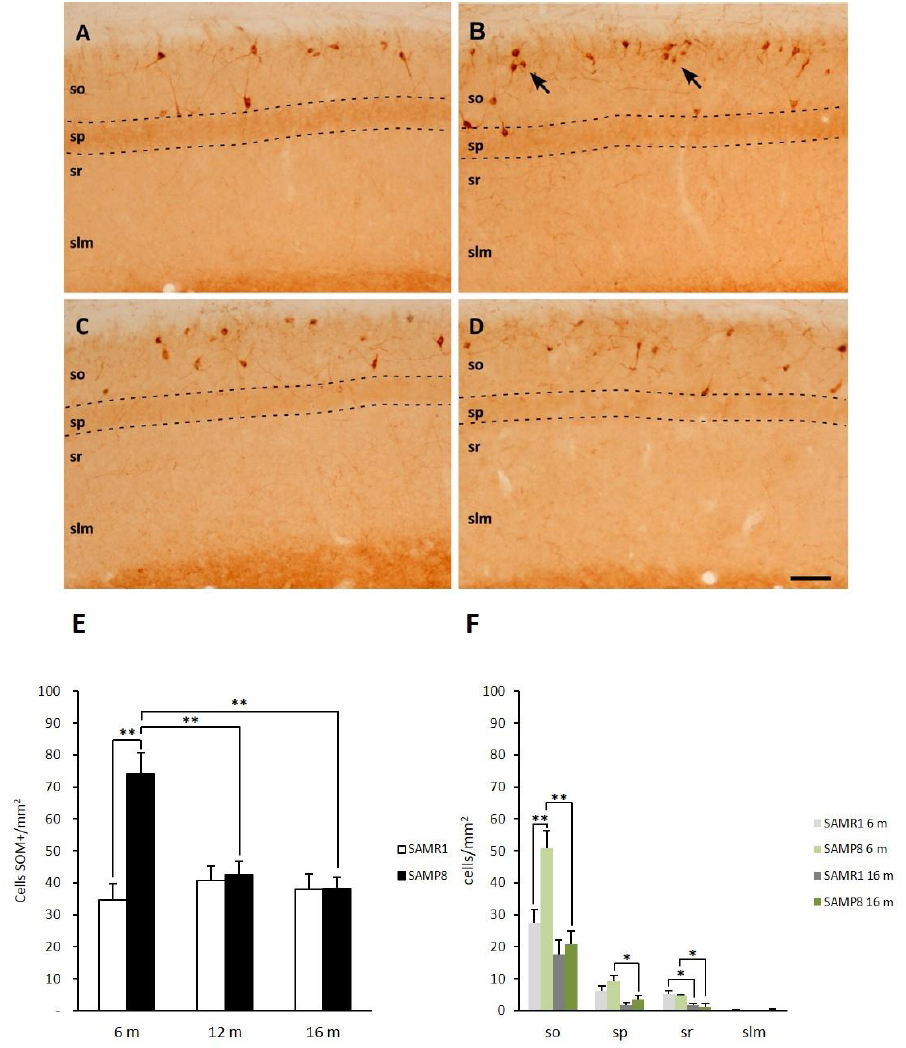
SST immunoreactive in CA1. By using ImageJ software and confocal picture we measured the number of SST neurons per mm^2^. Measurements in SAMR1 and SAMP8 mice were conducted at three different points of age: 6 months-old, 12 months-old and 16 months-old. A-D) DAB immunohistochemistry for SST(SOM) in 6months-old SAMR1(A), 16 months-old SAMR1(C), 6months-old SAMP8(B), and 16months-old SAMP8 mice(D). The pictures show CA1 field. There were SOM+ neurons accumulate in clusters in 6 month-old SAMP8 mice. E) Density of SST(SOM) neurons in the whole CA1 field. F)density of SST neurons in each of the CA1 subfields. *; p-value ≤0.05. **; p-value ≤0.01.. so; *stratum oriens*. sp: *stratum pyramidale*. sr: *stratum radiatum*. Slm: *stratum lacunosum moleculare*. Scale bar: 100µm.

When comparing SAMR1 and SAMP8 mice at 6, 12, and 16 months-old, a statistical difference was found at 6 months of age (p-value: 0.001), where the number of SST+ cells per area was a 40% larger in 6 months-old SAMP8 than 6 months-old SAMR1. (SAMR1: 34.63 ± 12.50 cells/mm^2^; SAMP8: 74.25 ± 15.56 cells/mm^2^). However, no statistical differences were observed at 12 and 16 months-old mice between SAMR1 group and SAMP8 group (12 months - SAMR1: 43.79 ± 10.77; SAMP8: 42.42 ± 10.63 cells/mm^2^; 16 months - SAMR1: 37.92 ± 11.86; SAMP8: 38.23 ± 8.49 cells/mm^2^).

Since we found changes of SST neurons in SAMP8 animals, a more detailed study of SOM interneurons was conducted and the distribution of these interneurons was analyzed in CA1 subfields. Therefore, the density of SST+neurons was independently measured in *stratum oriens, pyramidal, radiatum* and *lacunosum-moleculare*. In SAMR1, the highest density of the SOM neurons was found in *stratum oriens* which was close to 20-30 SST+ neurons/mm^2^. However, in the rest of subfields of CA1 this population did not reach 10 SOM+ neurons/mm^2^. Although there was not a significant decrease of SST in the whole CA1 region related to aging process of SAMR1, there was a tendency to decrease in all subfields and in deed, the density decrease between 6 months-old SAMR1 and 16 months-old SAMR1 mice was significant in *stratum radiatum* (p-value < 0.05).

Equally, the higest density of SST interneurons is located in *stratum oriens* of SAMP8 mice. However, different from what happened on SAMR1 mice, there is a significant decrease of SST interneurons in *stratum oriens, pyramidal* and *radiatum* during the aging process of SAMP8 mice. It is important to highlight that the population of SST interneurons decrease gently in *stratum pyramidal* and *radiatum* and there (6 months – SAMP8: 4.95 ± 1.85 cells/mm^2^. 16 months – SAMP8: 1.74 ± 1.71 cells/mm^2^) is a more sudden change in *stratum oriens* (6 months – SAMP8: 50.79 ± 10.77 cells/mm^2^; 16 months – SAMP8: 21.92 ± 9.85 cells/mm^2^).

Comparing 6 months-old SAMR1 and SAMP8 there is a clear different of the density of SST interneurons in the *stratum oriens*, which, is vanished at 16 months old of age. On the other hand, the other regions analyzed did not show intergroup differences. Therefore, the neuron population that was mainly changed in SAMP8 mice belongs to O-LM neurons, which are SST interneurons whose somas locate in *stratum oriens* and which depict more than 90% of the SST positive neurons in CA1. Since those interneurons have a specific target on dendrites of the principal neurons in the *stratum lacunosum-moleculare*, a quantitative analysis was carried out by densitometry analysis in the *stratum lacunosum-moleculare* of CA1. This study confirmed that the density of neuropil was statistically higher (p: 0.002) in 6 months-old SAMP8 than in 6 months-old SAMR1 (SAMR1: 3.44 ± 0.89; SAMP8: 4.33 ± 0.86 cells/mm2). However, in elder animals, the neuropil optic density was statistically lower (p-value: 0.001) in 16 months-old SAMP8 than in 16 months-old SAMR1 mice (SAMR1: 5.04 ± 0.74; SAMP8: 3.96 ± 0.66 cells/mm2). Additionally, the density of neuropil increased significantly between 6 months-old and 16 months-old SAMR1 mice (figure 3).

**Figure 3.**
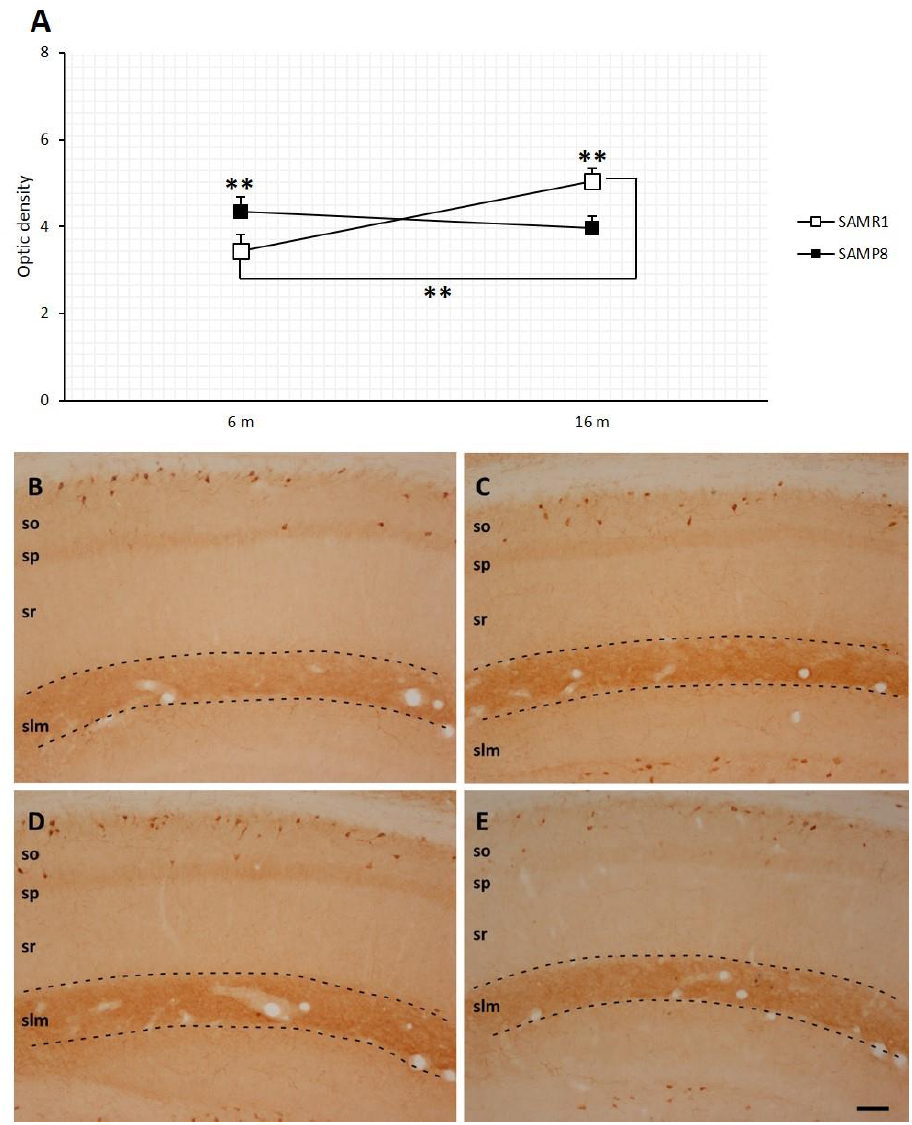
Optical density of SST+ labeling in the hippocampus of SAMR1 and SAMP8 mice. **A**: the graph depicts the changes of optical density between young and aged mice. B-E: representative pictures showing IHC against SST in 6 months-old SAMR1 (B), 6 months-old SAMP8 (C), 16 months-old SAMR1 (D), and 16 months-old SAMP8 (E). Dotted line delimit *stratum lacunosum-moleculare*. **; p-value ≤ 0.01. so; *stratum oriens*. sp: *stratum pyramidale*. sr: *stratum radiatum*. slm; *stratum lacunosum-moleculare*. Scale bar: 100 µm.

##### D. Cholinergic system

Since alterations of cholinergic system have been shown in the septum-hippocampal pathway of SAMP8 mice and because the O-LM neurons receive projection from the septum, we decided to analyze the immunoreactivity against choline acetyl-transferase (ChAT) as control for our study.

Densitometry analysis showed some differences between SAMP8 and SAMR1 mice. From a first qualitative analysis, these differences were already noticeable in 6 months-old SAMP8, which showed bigger values of optical density than 6 months-old SAMR1. Those differences were found both in the hippocampus (SAMR1: 0.94 ± 0.05; SAMP8: 1.14 ± 0.15) and the CA1 *lacunosum-moleculare* layer (SAMR1: 0.81 ± 0.08; SAMP8: 1.27 ± 0.2). Both hippocampus (p-value: 0.01) and *lacunosum-moleculare* (p-value: 0.002) differences were statistically significant. As in the case of SOM densitometry analysis, changes of plexus density were qualitatively appreciable in the *stratum lacunosum-moleculare* of CA1 of all SAMP8 studied animals as well(figure 3). In the present case, the 16 months-old SAMP8 had higher density of neuropil ChAT immunoreactive than 6 months-old SAMP8 mice. Interestingly, we observed similar behavior in SAMR1, where the immunoreactivity against ChAT is higher in older than in younger SAMR1 mice (figure 4).

**Figure 4.**
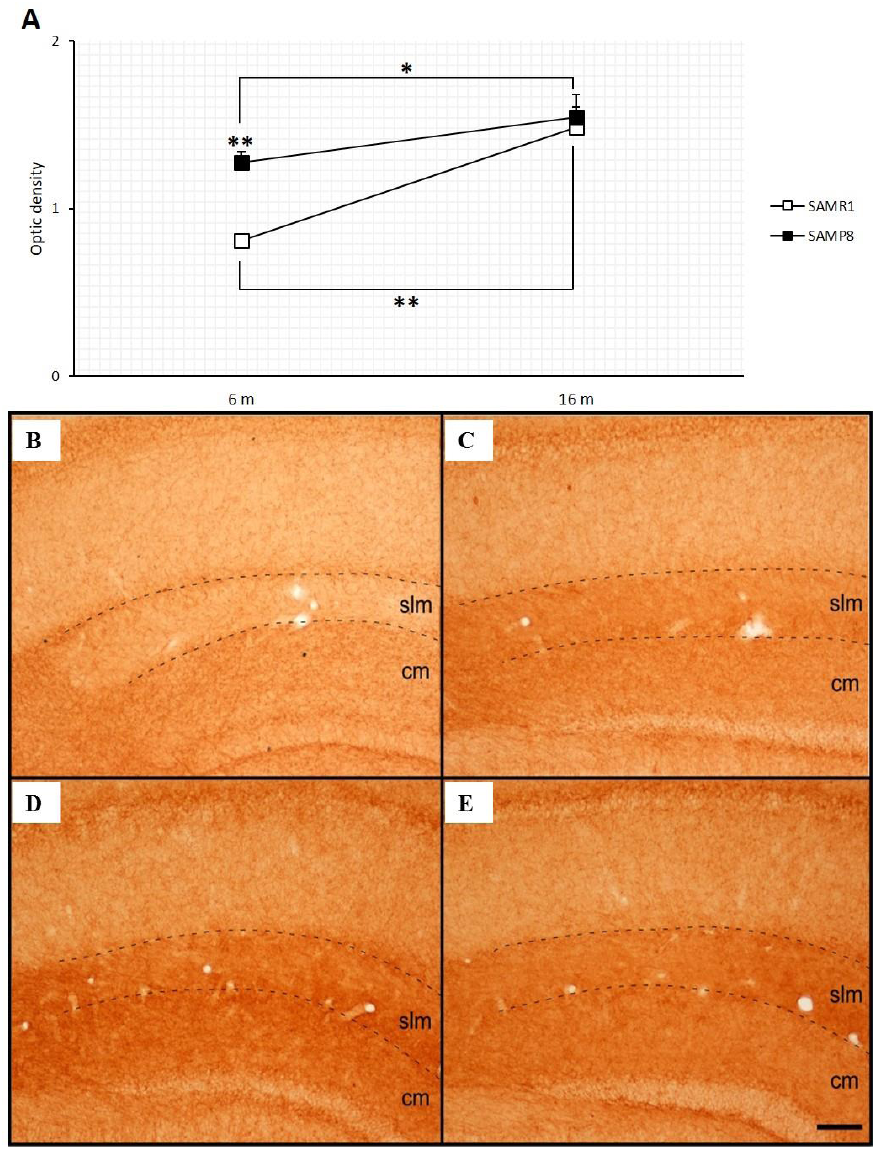
A: Optical density of ChAT labeling in the CA1 fields of SAMR1 and SAMP8 (n=16). B-E: microphotographs of DAB immunohistochemistry for ChAT in CA1. The images show different patterns of labeling between the 6 months-old SAMR1 and SAMP8. 6 months-old SAMP8 mice always showed a stronger signal in the *stratum lacunosum-moleculare* than SAMR1. In older mice, this difference disappears. B: 6 months-old SAMR1; C: 6 months-old SAMP8; D: 16 months-old SAMR1; E: 16 months-old SAMP8. slm: *stratum lacunosum-moleculare*; ml: molecular layer of dentate gyrus. Scale bar: 100μm. E; average optical density of the ChAT in the *stratum lacunosum-moleculare* signal at 6 and 16 months.

##### E. Double labeling: CR and SST immunopositive interneurons

Double immunofluorescence for CR and SST was conducted in order to assess whether there was any change in the expression of this neuro markers in SAMP8 mice or not. Data shown there was not difference between SAMR1 and SAMP8. In like manner, the number of double CR/SST+ neurons per mm^2^ at 6 months of age was similar in both groups and equally, the densities of double CR/ SST+ neurons of 16 months-old SAMR1 and 16 months-old SAMP8 mice had similar values. However, a significant decrease of this neural population was observed when 6 months old mice and 16 months old mice of both SAMR1 and SAMP8 groups were compared. The results showed that CR+/SST+ neurons lost is greater than the loss of SST+ or CR+ neurons during the aging process (figure 5E).

**Figure 5.**
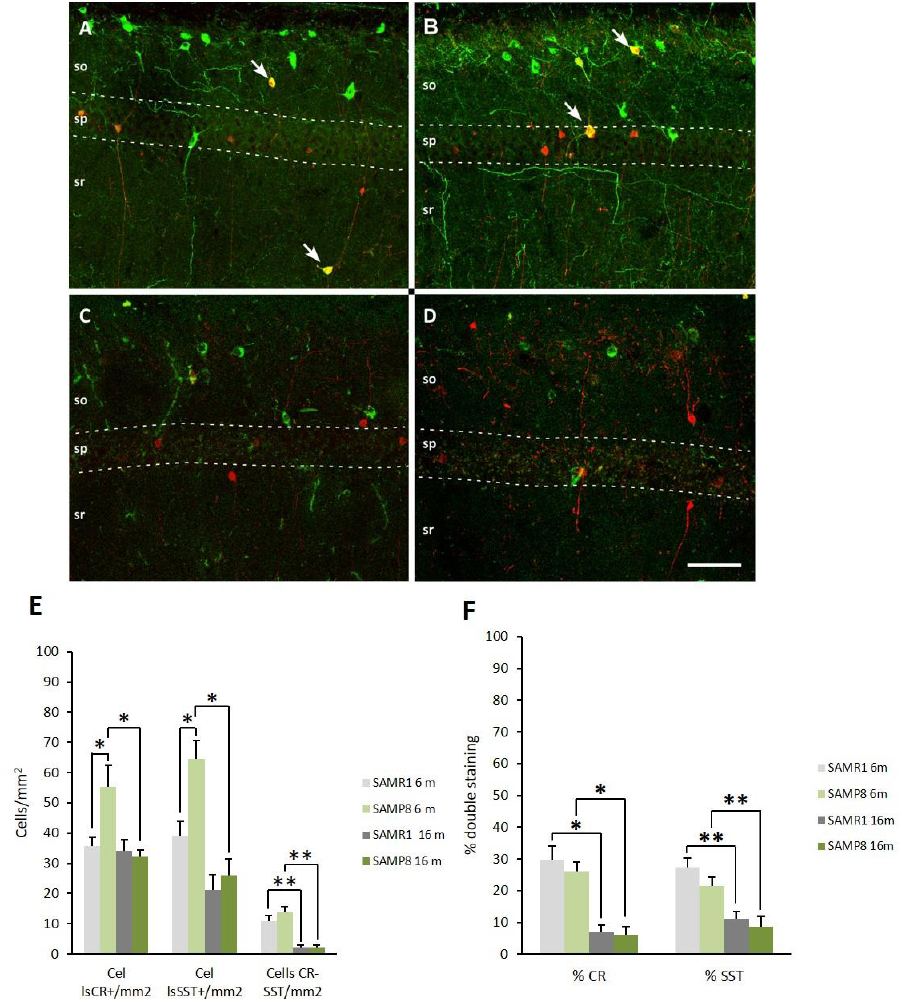
Double staining against CR and SST. Assessment and comparison of the density of CR+, SST+ and CR+/SST+ neurons in CA1. A-D: Double immunostaining against CR and SST in 6 months-old SAMR1 (A), 6 months-old SAMP8 (B), 16 months-old SAMR1 (C), and 16 months-old SAMP8 (D) mice. In the younger mice, CR+/SST+ neurons were found more often than in the elder animals (arrows). Scale bar: 100 µm. E: the graph depicts the density of CR+ neurons, SST+ neurons and CR+/SST+ neurons. F: percentage of CR+ and SST+ neurons that are double stained. *: p-value ≤ 0.05. **: p-value ≤ 0.01. so: *stratum oriens*. sp: *stratum pyramidale*. sr: *stratum radiatum*. slm: *stratum lacunosum-moleculare*. Scale bar: 100 µm.

In order to be able to determine if this population of CR and SST+ neurons declined in a proportional manner to the observed decrease in the populations of CR and SST+ interneurons we calculated the percentage of CR+ neurons that are SST + (CR% in figure 5F)and the percentage of SST+ neurons that are CR+ (SST% in figure 5F). This data showed that this population of double is more affected by the aging process than other SST+ and CR+ subset of neurons.

## DISCUSSION

### Hipocampal volume along aging

In order to discard that any of the density changes where due to changes in the hippocampal volume, we measure the volume of the hippocampi at the three points of age. SAMR1 mice did not showed changes of hippocampus volume between the 6 months-old SAMR1 and 16 months-old SAMR1.Thus there was no changes of the hippocampal volume related to the aging process. Although there was not clear change in the volume of the hippocampus of SAMP8, unlike SAMR1, the hippocampi of SAMP8 tend to decrease with the aging process (p-valor = 0.06). This data it is in line with some previous studies where the decrease of hippocampal volume has been found in neurodegenerative diseases (de Jong, van der Hiele et al. 2008, Bangen, Preis et al. 2017, Tangaro, Fanizzi et al. 2017).

### CR+ and SST+ interneurons of SAMP8 change during aging compared to controls

The densities of CR+, SST+ and PV+ neurons did not significantly change in CA1 field of SAMR1 between 6 and 16 months old of age. Contrary, CR + and SST + interneurons decrease significantly between young and aged SAMP8 mice.

Interestingly, the population of CR+ and SST+ interneurons was larger in 6 months-old SAMP8 mice than in 6 months-old SAMR1, but no significant differences were found comparing 16 months-old SAMR1 and 16 months-old SAMP8 mice.

Most of the CR-immunoreactive neurons of CA1 belong to the kind of IS-interneurons, which are interneurons that innervate other interneurons (Baude, Nusser et al. 1993, Ferraguti, Cobden et al. 2004, Chamberland, Salesse et al. 2010), and likely, this set of interneurons is increased in 6 months-old SAMP8 mice. However, futher studies are needed to support this hypothesis.

On the other hand, the population of SST + interneurons is strongly altered in SAMP8. At 6 months-old SAMP8 mice have larger density of SST+ neurons than 6 months-old SAMR1. In CA1 field, SST + cells are almost entirely restricted to *stratum oriens* and belong to a specific population of interneurons called O-LM (Gulyas, Hajos et al. 2003, Stanley, Fadel et al. 2012). Therefore, this population of interneurons is changed in SAMP8 mice.

Several behaviour tests have shown that learning and memory impairments already exist in 4 and 6 months-old SAMP8 mice(Miyamoto, Kiyota et al. 1986, Miyamoto 1997) and the onset of synaptic connections loss and development of beta-amyloid plaques in SAMP8 occur around this age as well(Del Valle, Duran-Vilaregut et al. 2010). We have found changes in the density of the hippocampal neurons around this early stage of 6 months-old, so it might be a link between the cognitive impairments found in SAMP8 and our data. Furthermore, the beginning of the accumulation of beta-amyloid plaques has been associated with a temporary increase in the expression of SST, followed by a subsequent reduction of this neurons (Geci, How et al. 2007, Tallent 2007). That might explain the increase in the density of SST+ neuron population in 6 months-old SAMP8 mice and the drastic decrease between 6 months and 12 months-old. For our future studies, it might be relevant to analyse how these populations of neurons and beta-amyloid plaques are interconnected from earlier stage of postnatal development and they relationship with the behaviour performance during hippocampal dependent tasks.

### Compensatory systems operating in CA1 of SAMR1 and SAMP8 mice

Since cholinergic system is involved in cognitive functions (Everitt and Robbins 1997, Hasselmo and Stern 2006) and, impairments of the cholinergic system has been already reported in the brains of SAMP8, we decided to study the immunoreactivity of ChAT in CA1 field in order to give more information about in which specific way the cholinergic system might be affected.

There are evidences, which report that SAMP8 mice have changes in the cholinergic system from the 4 months of age. It has previously been reported that the activity of ChAT enzyme in the brains of SAMP8, between 4 months and 12 months old, decrease up to 50% (Strong, Reddy et al. 2003) and, there is an overexpression of butyrylcholinesterase enzyme, (an enzyme involved in catalyzing the hydrolysis of acetylcholine in the synaptic clefts) in the brains of 4 months-old SAMP8 animals (Fernandez-Gomez, Munoz-Delgado et al. 2008). Following this line, our results showed a higher optical density of ChAT in the stratum *lacunosum-moleculare* of CA1 in 6 months-old SAMP8 mice compared to their controls, and that might be related to an accumulation of the soluble form of enzyme in the synaptic terminals of *lacunosum-moleculare* layer. Mostly the septo-hippocampal cholinergic projections target this region of the hippocampus and the cholinergic neurons will synapse onto the dendrites of CA1 principal cells (Teles-Grilo Ruivo and Mellor 2013). Supporting our data, an increase of ChAT activity has been reported in the hippocampus and the frontal cortex (DeKosky, Ikonomovic et al. 2002, Counts and Mufson 2005)of patients with mild cognitive impairments (MCI, a prodromal stage of Alzheimer disease). In addition, a disconnection of glutamatergic entorhinal cortex input to the hippocampus occurs early in the Alzheimer disease (Gomez-Isla, Price et al. 1996, Kordower, Chu et al. 2001). Hence, former results support the hypothesis that the upregulation of hippocampal ChAT in MCI cases may be due to the replacement of enervated glutamatergic synapses by cholinergic input arising from the septum(Mufson, Counts et al. 2008). This is emphasized by animal experimental studies demonstrating reactive sprouting in the hippocampus following perforant path transsections in rats(Savaskan and Nitsch 2001). However, this increase in ChAT activity is not observed in Alzheimer cases, so it might be an early sympthom related with an increased risk of developing Alzheimer’s disease(Schliebs and Arendt 2011). This might explain why 6 months-old SAMP8 mice have a higher optic density in the *stratum lacunosum-moleculare* compare to their controls and this difference disappear in aged mice. Likely the higher density of SST+ neurons in 6 months-old SAMP8 mice is linked to the cholinergic system since septal cholinergic neurons target O-LM interneurons as well(Teles-Grilo Ruivo and Mellor 2013). Moreover, axons of O-LM neurons project to *lacunosum-moleculare* where septal cholinergic neurons are stimulating principal neurons. It might exist a compensatory mechanism between inhibitory and excitatory system in this area of the hippocampus. Additionally, our data show that there is a significant increase of optical density of SST and ChAT immunoreactivity in 16 months-old SAMR1 mice, therefore this compensatory mechanism might occur during normal aging as well. However, later in time.

Interestingly, although there is an increase of CR and SST neuron populations in 6 months-old SAMP8 mice compare to its control, the ratio of SST/CR+ neurons in SAMP8 mice is similar to SAMR1. Equally, 16 months-old SAMR1 and 16 months-old SAMP8 mice have similar ratio of SST/CR+ neurons in CA1 despite the decrease of these neurons during the aging process (supplementary figure 3). Thus, the same compensatory systems of local and long-range projections have a key role in pathological and normal aging. More detailed studies of how each of these set of populations affect the local network of the hippocampus can help to understand the onset of neurodegenerative disease and the different to normal aging.

### Remarks

Further electrophysiology and gen regulation studies on this local network might be relevant in order to understand the reason of why these populations are increased in young adults and decreased so drastically during the aging process. Additionally, this direction of research can help to give new hints about how neurodegenerative diseases develop as well as how interneuron populations are regulated and affect to the the local network.

Most of the changes that we have reported in this study are tightly related to the *stratum-lacunosum moleculare*, where CA1 receives its main input from entorhinal cortex through the perforant pathway. More detailed studies of the parahippocampal inputs into the hipocampus may be needed. Neuroantomical studies of the perforant pathway might help to understand the origin of the neurodegenerative diseases. It is quite accepted that parahippocampal region is clearly damage in early stages of several neurodegenerative diseases. So, finding if changes of entorhinal cortex are first in time than hippocampal changes or not, and finding which specific neurons are targets of these impairments might become one of the main challenges of this century. In the current paper, we propose SAMPR1/P8 model as candidates to help in answering these questions.

## SUPPLEMENTARY FIGURES

**Supplementary figure 1.**
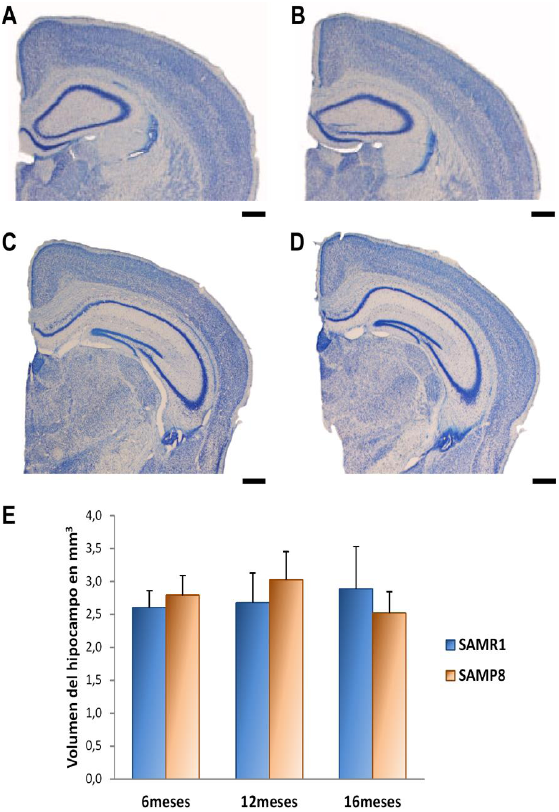
Hippocampal volume measured in SAMR1 and SAMP8 animals. Nissl staining of 6 months-old SAMR1 (A), 6 months-old SAMP8 (B), 16 months-old SAMR1 (C) and 16 months-old SAMP8 (D). The pictures show examples of the more rostral and caudal levels (Bregma −1.22 and Bregma −2.54 respectively) that we chose for the hippocampal volume measurement along the rostral–caudal axis. Scale bar: 500 µm. E: averages of the hippocampal volumes at three age points. There were no significant changes in hippocampal volume associated with aging.

**Supplementary figure 2.**
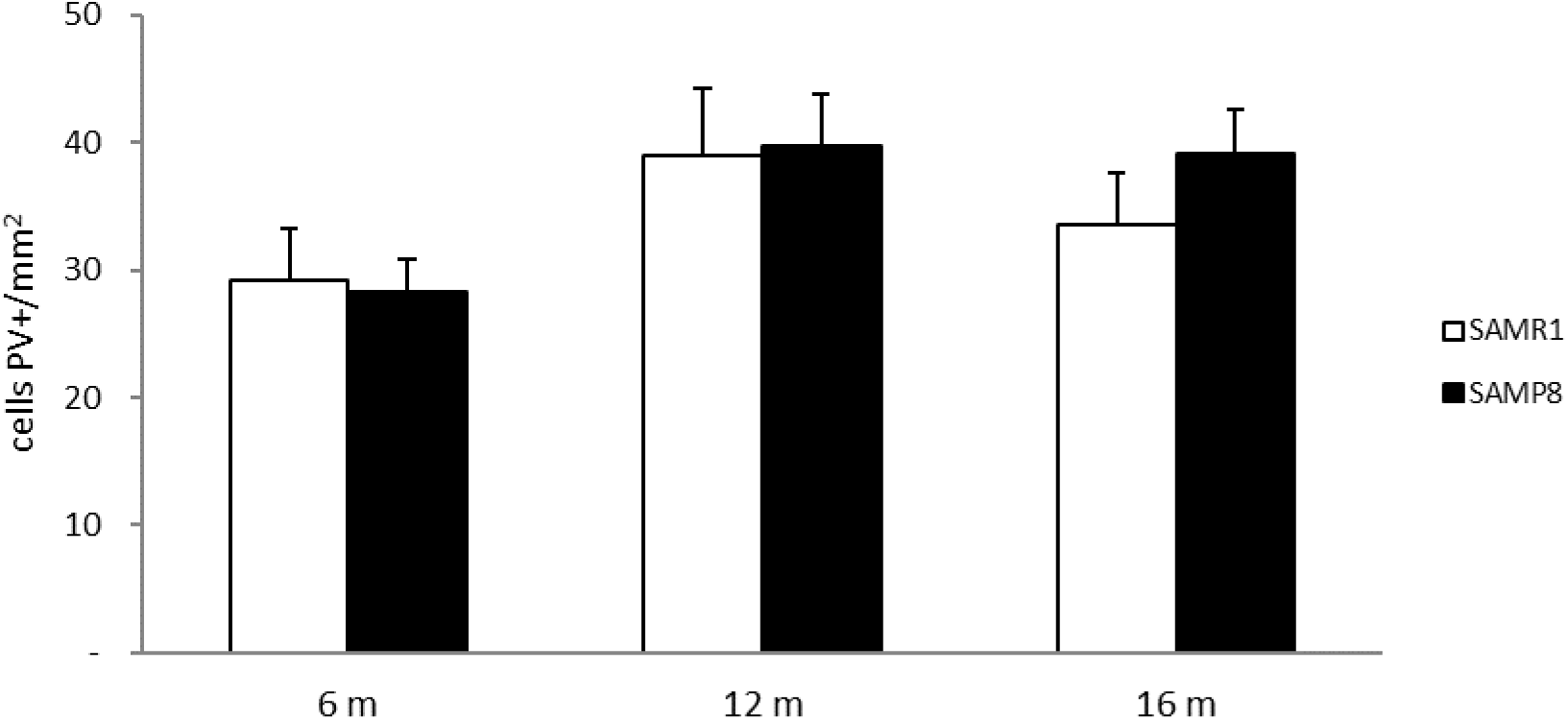
Density of the population of PV+ cells in CA1. The graph shows the average densities of the PV+ interneurons in CA1 of SAMR1 and SAMP8 mice at 6, 12, and 16 months of age. There were not significant differences between groups.

**Supplementary figure 3.**
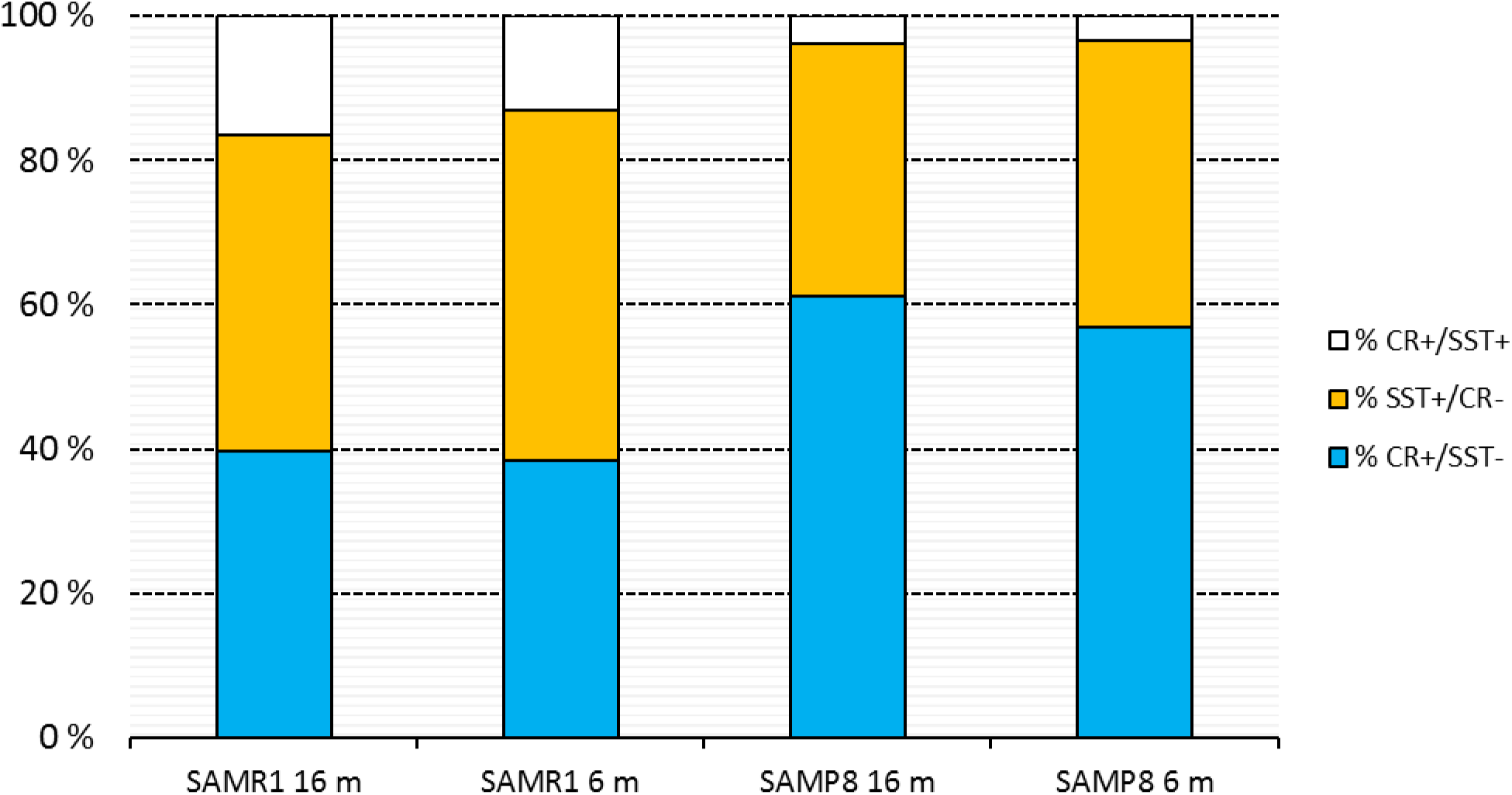
Percentage of neurons that are SST+, CR+ and SST+/ CR+ in CA1. The percentage of each neuronal type is not affected by aging, but differences between SAMR1 and SAMP8 groups can be observed.

